# Wolves in black: multiple introgressions and natural selection may explain melanism in Italian wolves

**DOI:** 10.64898/2026.05.08.723698

**Authors:** Fabbri Giulia, Battilani Daniele, Mattucci Federica, Galaverni Marco, Stronen Astrid Vik, Musiani Marco, Godinho Raquel, Lobo Diana, Scandura Massimo, Randi Ettore, Fabbri Elena, Caniglia Romolo

## Abstract

Hybridisation between wild and domestic taxa can favour the spread of domestic alleles into wild populations through backcrossing. The complex interplay of random genetic drift, recombination, and selection can shape the fate of introgressed alleles. Maladaptive domestic variants are likely to be purged by natural selection, but others may persist across generations. It has long been known that the Apennine Italian wolf population, exposed to large numbers of free-ranging dogs, has experienced extensive introgression. The unusually high frequency of black wolves observed in Italy, compared to other European populations, may parallel patterns documented in North American wolves, where the melanistic *K^B^* allele at the CBD103 gene, of domestic origin, has spread over thousands of years of introgression. We tested whether the *K^B^* mutation entered the peninsular Italian wolf population via hybridisation and spread through adaptive introgression. Genome-wide analyses of black and wild-type (grey-coated) Apennine wolves showed no clear signatures of recent dog ancestry in most melanistic animals. Our ancestry reconstruction approaches identified two distinct *K^B^* haplogroups of domestic origin, suggesting multiple introgression events. Notably, we found molecular evidence consistent with balancing selection on the *K^B^* haplotypes, whose functional role, nonetheless, warrants further research. Therefore, the microevolutionary genomic and ecological consequences of wolf-dog hybridisation in Italy should be carefully investigated to inform appropriate science-based conservation management strategies.

## Introduction

The grey wolf (*Canis lupus,* Linnaeus 1758) was the first species to be domesticated, at least 30,000 years ago (vonHoldt et al. 2010; Thalmann et al. 2013; Shannon et al. 2015; Frantz et al. 2016; Bergström et al. 2022). During this process, domesticated proto-dogs and dogs (*Canis lupus familiaris*) underwent both unintentional and directional selective pressures to develop human-desirable behavioural and morphological traits such as boldness, tameness, coat colour, body size, and shape (Cadieu et al. 2009; Boyko et al. 2010; Banlaki et al. 2017; Pendleton et al. 2018). However, despite thousands of years of domestication, dogs and wolves have remained interfertile and have crossbred, producing fertile offspring (Vilà and Wayne 2001; Godinho et al. 2011; Fan et al. 2016; Galaverni et al. 2017; Pilot et al. 2018; Lin et al. 2025). Paleogenomic studies have shown recurrent bidirectional gene flow between wolves and dogs since the initial stages of domestication (Freedman et al. 2014; Ciucani et al. 2019; Bergström et al. 2022). Over the past few decades, genetic and genomic tools have identified cases of recent wolf-dog hybridisation in several European wolf populations (Dziech 2021). Evidence of prolonged canine introgression into wolf genomes, observed mainly at local scales in certain Mediterranean countries, has raised important evolutionary and conservation issues (Randi 2008; Freedman et al. 2014; Larson and Fuller 2014; Donfrancesco et al. 2019; Bergström et al. 2020; Battilani et al. 2025). Indeed, the introgression of exogenous alleles into wild populations is expected to change allele frequencies and affect the direct or pleiotropic expression of phenotypes, including at adaptive loci such as those related to life history traits and reproductive fitness (Ducrest et al. 2008; Todesco et al. 2016; Pilot et al. 2021; Schneider et al. 2025). Although the outcomes of wolf-dog hybridisation and introgression are specific to each population (Hedrick 2013), they are generally assumed to have negative effects on fitness and survival in the wild because many domestic traits have evolved under relaxed or divergent selective regimes compared to natural conditions (Bjornerfeldt et al. 2006; Rubin et al. 2012; Theodoropoulos et al. 2025). However, empirical support for these effects is still lacking.

According to population genetic theory, random drift might prevail over weak purging selection pressures in small, isolated, and fragmented populations (Hedrick 2013), as repeatedly demonstrated by empirical genomic evidence for wolves in Western Europe (Pilot et al. 2021; Battilani et al. 2025).

However, some domestic variants that have entered the gene pool of wild populations can confer adaptive advantages (Hindrikson and Tammeleht 2025; Lobo et al. 2025; Sarabia et al. 2025), as seen in the case of domestic goat (*Capra hircus*,) MHC haplotypes introgressed into the genome of the Alpine ibex (*Capra ibex*), and domestic cat (*Felis catus*) immune-related alleles introgressed into the genome of the Scottish wildcat (*Felis silvestris*) (Grossen et al. 2014; Howard-McCombe et al 2023). Another notable example of adaptive introgression is a melanistic mutation at the *β*-defensin gene *CBD103*, which most likely originated in dogs and spread into North American and Italian wolves, coyotes (*C. latrans*) and golden jackals (*C. aureus*; Anderson et al. 2009; Galov et al. 2015).

Coat colour in dogs is controlled by complex epistatic interactions between the Agouti signalling protein (*ASIP* locus) and the melanocortin-1 receptor (*MC1R*), and at least 12 other genes (Schmutz et al. 2002; Kerns et al. 2004; Bannasch et al. 2021; Brancalion et al. 2021). Black coats in some dog breeds and in melanistic wolves are, however, mainly controlled by the so-called K-locus, corresponding to the *β*-defensin gene *CBD103* mapping to canine chromosome 16 (Candille et al. 2007; Anderson et al. 2009). The molecular mechanism involves the transmembrane protein encoded by the MC1R receptor, and two alternative ligands: the *ASIP* protein and the peptide encoded by *CBD103*. The product of the dominant melanistic *CBD103 K^B^*allele, a three-nucleotide deletion, interacts with *MC1R* and prevents *ASIP* from binding, thereby stimulating the production of eumelanin (black or brown pigment). In contrast, the recessive wild-type *k^y^* allele does not affect the interaction between *MC1R* and *ASIP*, leading to the production of brownish pheomelanin (yellow to nearly white pigment). However, not all melanistic canids carry the *K^B^* allele since other genes are involved in the pathway, including *ASIP* (Kerns et al. 2004; Bannasch et al. 2021) and *TYRP1* (Schmutz et al. 2002). In recent years, both the genetic basis and the origin and potential fitness advantages of melanism in North American grey wolves have been clarified (Anderson et al. 2009; Coulson et al. 2011; Stahler et al. 2013; Cassidy et al. 2017). Schweizer et al. (2018) showed that the *K^B^* allele was probably introgressed from early Native American dogs into Canadian Yukon wolves in the last 7,000 years and rapidly spread in North American wolves due to its adaptive potential. North American wolf populations exhibit a clear latitudinal cline in coat colour, with pale phenotypes predominating in the High Arctic and black phenotypes in forested regions (Gipson et al. 2002; Musiani et al. 2007). Although camouflage or thermoregulation may contribute to this pattern, the process regulating the spread of the black coat appears more complex and related to the immunological role of the β-defensin genes (Anderson et al. 2009; Coulson et al. 2011; Hedrick et al. 2016; Johnston et al. 2021). Indeed, low-frequency introgressed genotypes that confer fitness advantages might maintain genetic polymorphisms in populations through assortative mating and balancing selection (Hedrick 2007).

Cubaynes et al. (2022) found that canine distemper virus (CDV) outbreaks can generate heterozygote advantage through fluctuating selection affecting the *K^B^* allele frequency and favouring disassortative mating behaviours, as reported for the case study of the black wolves in the Yellowstone National Park (Hedrick et al. 2014; 2016). Despite widespread wolf-dog hybridization, black wolves have rarely been reported on other continents (Apollonio et al. 2004; Godinho et al. 2011; Doan et al. 2023; Roda and Philibert 2025). A notable exception is found in Italy, where black individuals have been frequently observed since the 1970s (Boitani 1983; Apollonio et al. 2004; Caniglia et al. 2013). The presence of the *K^B^* allele in Italian wolves (*C. l. italicus*) has been genetically monitored over the last 15 years by routinely analysing DNA from invasively and non-invasively collected biological samples, allowing its distribution to be assessed at a national scale (Randi et al. 2014; Caniglia et al. 2020; Gervasi et al. 2024). However, despite the documented recurrent CDV presence in wild mammals (Di Sabatino et al. 2014, 2016; Bianco et al. 2020; Ricci et al. 2021), it remains unclear whether selective pressures like those acting in North American grey wolves might explain the spread of the melanistic phenotype in Italian wolves. Galaverni et al. (2017) found a strong association between the black coat and single nucleotide polymorphisms (SNPs) surrounding the K-locus, and confirmed the domestic origin of this region through local ancestry inference and haplotype reconstruction. However, the timing of its introgression into Italian wolves remains unresolved, and multiple admixed origins have been hypothesised (Galaverni et al. 2017).

To further clarify these questions, we analysed the genetic profiles of a set of Italian phenotypically black and wild-type (grey-coated) wolves at 170k SNPs (Galaverni et al. 2017) and compared them with corresponding data from North American wolves, village dogs, and golden jackals. We used population genomic tools to obtain genome-wide ancestry reconstructions and evidence of haplotype selection in the black Italian wolves. Results were evaluated to test whether: 1) the melanistic *K^B^* deletion in the Italian wolf population originated from a single domestic dog introgression or from multiple events; and 2) the diffusion dynamics of the domestic *K^B^* haplotypes into the Italian wolf population have been shaped by random genetic drift or by some form of selective pressure.

## Materials and methods

### Dataset collection and genotype filtering

The dataset we generated included all unrelated individuals: 14 wild-type Italian wolves (seven males, seven females), 14 black Italian wolves (six males, eight females), and 28 Italian village dogs (14 males, 14 females). Additionally, we added three wild-type (one male, two females) and four black (one male, three females) North American wolves for comparative purposes, and 18 (nine males, nine females) wild-type golden jackals as outgroup species (**Table 1**). Samples were genotyped with the Illumina Canine HD Beadchip with 170K SNPs, which were mapped on the CanFam2 version of the dog genome. The presence/absence of the 3-bp deletion at the CBD103 gene was diagnosed by fragment amplification using specifically designed and optimized primers (Caniglia et al. 2013). We selected autosomal SNPs with no more than 5% missing data and minor allele frequency (MAF) > 0.05 (hereafter: quality-pruned dataset) in PLINK v1.90b6.26 (Purcell et al. 2007), and further pruned for linkage disequilibrium (LD) with the option --indep-pairwise 50 5 0.2 (LD-pruned data set).

**Table 1:** Dataset used in the present study. Acronym, number of analysed samples, and sampling period are given per group.

| Acronym | Group | N | Presence of $K^B$ allele | Sampling period |
| --- | --- | --- | --- | --- |
| WIW | Wild-type Italian wolves | 14 | No | 1991-2015 |
| BIW | Black Italian wolves | 14 | Yes | 2000-2014 |
| WNAW | Wild-type North American wolves | 3 | No | 1999-2002 |
| BNAW | Black North American wolves | 4 | Yes | 1999-2000 |
| ID | Italian dogs | 14 | No | 2005-2014 |
| BID | Black Italian dogs | 14 | Yes | 2005-2014 |
| WGJ | Wild-type East-European golden jackals | 18 | No | 2009-2013 |

### Population structure

Population structure was investigated on the LD-pruned dataset through principal component analysis (PCA) and a maximum likelihood model-based ancestry procedure, using ADMIXTURE (Alexander et al. 2009), applying both methods to the genome-wide set of markers and to SNPs restricted to chromosome 16 only (hosting the *CBD103* gene). PCA was performed in R with the package ‘adegenet’ v2.1.3 (Jombart 2008). ADMIXTURE v1.22 (Alexander et al. 2009) was used to determine the most likely number of ancestral components (K) in the dataset between K = 2 to K = 6, based on the lowest cross-validation error, and to describe each individual’s genetic make-up according to its genome-wide ancestry proportions upon variation of the number of K.

### Admixture and local ancestry inferences

We formally tested for wolf-dog gene flow in the Italian wolves carrying the *K^B^* allele using the D-statistics implemented in the R package ‘ADMIXTOOLS2’ (Maier et al. 2023) on the quality-pruned dataset. First, we tested whether BIW had an excess of allele sharing with dogs (ID or BID) compared to WIW. The D-statistics analysis was carried out genome-wide and on chromosome 16 only, testing two different topologies in the form of (((P1, P2), P3), P4): (i) gene flow between each Italian wolf individual (X) or each other Italian wolf individual (IW) relative to BID using WGJ as the outgroup (((X, IW), BID), WGJ); (ii) gene flow between each Italian wolf individual (X) or each other Italian wolf individual (IW) relative to ID using WGJ as the outgroup (((X, IW), ID), WGJ). We also tested the same topologies on each North American wolf individual. No filters were applied to the genotypes as the software internally considers only non-missing SNPs in all four groups.

We implemented local ancestry inference (LAI) analyses on the BIW samples to visualize genome-wide introgression patterns and focus on locus-specific ancestry at the K-locus, leveraging the quality-pruned dataset. We first jointly phased all the Italian wolf and dog genotypes with SHAPEIT v2.837 (Delaneau et al. 2012) using the recombination map from Muñoz-Fuentes et al. (2015). Then, to establish the most likely ancestor for each haplotype, we used the machine learning approach (Random Forests) implemented in RFMix (Maples et al. 2013), which is very fast and robust to slight deviations from the real admixture time. We provided WIW and BID as source populations and ran RFMix with default parameters except for the admixture time that was set to 4 generations ago based on previous results (Galaverni et al. 2017).

The time since admixture for each BIW sample was estimated by applying the formula developed by Johnson et al. (2011): T=B/(2×2×0.01)×L×z(1-z) to RFMix results, where B is the expected number of ancestry switches between dog and wolf genomic blocks, L is the genome length in Morgans, and z is the genome-wide ancestry proportion from a parental (dog or wolf) population. The number of generations since the admixture was converted into years, assuming a wolf generation time of 3 years (Skoglund et al. 2011), which would be more appropriate for expanding populations (Wikenros et al. 2021) characterized by high human-related mortality potentially affecting local pack social structure and individual age of reproduction (Musto et al. 2021).

Furthermore, we used the R package ‘rehh’ (Gautier et al. 2017) to visualize the haplotype structure in BIW, considering the last 200 SNPs before the end of chromosome 16 encompassing the K-locus, obtained from the quality-pruned dataset. As our genotypes were obtained from a commercial SNPchip, they did not directly include information on the 3-bp deletion that characterizes the *K^B^*allele, so we manually added the presence/absence of the *K^B^* allele as a biallelic SNP with a central position with respect to the deletion.

### Selection analyses

We compared patterns of extended haplotype homozygosity (EHH) in the *k^y^*/*K^B^* pairs of Italian wolves, calculating allele-specific EHH in BIW around the K-locus. The ancestral and derived alleles were determined as the absence and presence of the 3-bp deletion for the K-locus, respectively. We specifically looked for selection signatures along chromosome 16 within each identified haplogroup in the BIW sample (see Results) as this method, being haplotype-based, would have no power in case of pooling together two or more different patterns of variation around the focus allele. Finally, based on previous evidence suggesting balancing selection at the K-locus in North American wolves (Schweizer et al. 2018), we explored whether a similar selective regime could also be operating in Italian wolves. Accordingly, we computed Tajima’s D with VCFTools (Danecek et al. 2011) in non-overlapping 200kbp windows (6 SNPs/window on average) across chromosome 16 in the separate BIW haplogroups, and we compared the obtained patterns with that in the WIW sample.

## Results

After filtering for missing alleles and MAF, the quality-filtered dataset consisted of 126K autosomal SNPs. After filtering for LD, the final LD-pruned dataset was composed of 14K SNPs and 438 SNPs for the genome-wide and chromosome 16 datasets, respectively.

### Population differentiation

In the genome-wide PCA analysis, the first principal component (PC1: 14.42%) primarily separated wolves and dogs from golden jackals, whereas the second component (PC2: 11.20%) differentiated Italian wolves (including both those carrying and not carrying the *K^B^* allele) from the other canids (**Figure 1A**).

**Figure 1:**
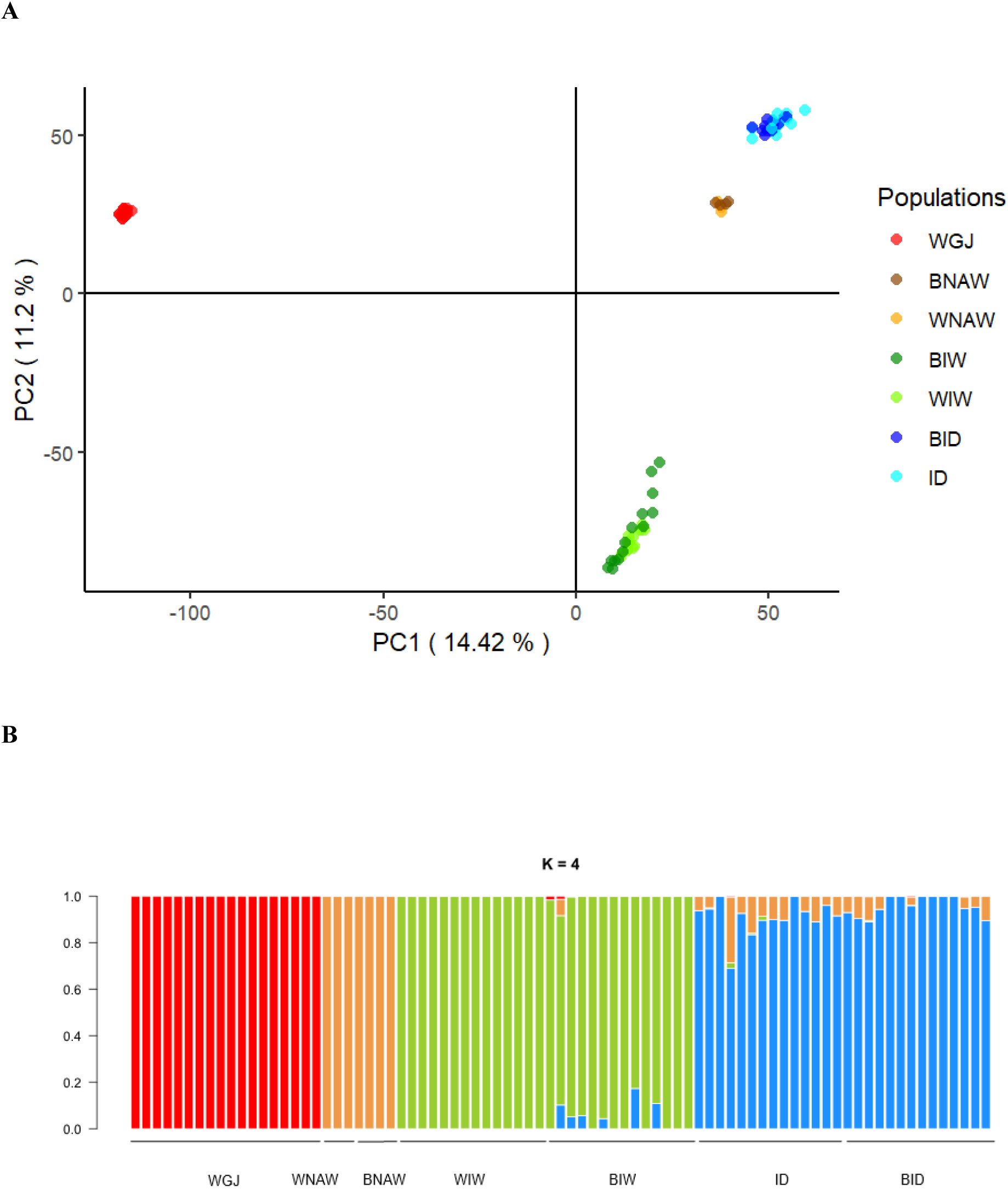
Population structure of the whole genome-wide LD-pruned dataset. (A) Principal component analysis (PCA) and (B) ADMIXTURE bar plot (K=4). WGJ = Wild-type East-European golden jackals; WNAW = Wild-type North American wolves; BNAW = Black North American wolves; WIW = Wild-type Italian wolves; BIW = Black Italian wolves; ID = Italian dogs; BID = Black Italian dogs.

ADMIXTURE analysis corroborated the PCA results. At K = 3 (**Supplementary Figure S1**), corresponding to the most supported number of populations according to the cross-validation error (**Supplementary Figure S2**), golden jackals, Italian wolves, and dogs clearly separated (mean estimated membership to the assigned clusters *Q1* > 0.999, *Q2* = 0.978, *Q3* = 0.980, respectively), while North American wolves apparently showed admixed ancestry and were mostly assigned to the dog cluster (mean estimated membership to cluster *Q3* = 0.652).

For this reason, we opted to visualize K = 4 (**Figure 1B**), which was consistent with the phylogenetic and geographic subdivision of the samples and indeed separated the North American wolf cluster as well, clarifying the ancestry assignment (**Supplementary Table S1**). In particular, golden jackals were included in cluster 1 (mean ancestry assigned to *Q1* > 0.999), North American wolves in cluster 2 (mean ancestry assigned to *Q2* > 0.999), Italian wolves in cluster 3 (mean ancestry assigned to *Q3* = 0.977), and dogs in cluster 4 (mean ancestry assigned to *Q4* = 0.933). When focusing on Italian wolves, all WIW and 8 BIW were entirely assigned to the wolf cluster 3 without dog ancestry (mean *Q3* > 0.999). The remaining 6 BIW showed an average domestic ancestry proportion of 9% with individual *q3* ranging from 0.814 to 0.955 (**Figure 1B; Supplementary Table S1**). Neither North American wolves nor golden jackals exhibited traces of dog ancestry (**Figure 1B; Supplementary Table S1**).

### Domestic introgression into the Italian wolf population

D-statistics results indicated that Italian wolves carrying the *K^B^* allele had an excess of allele sharing with dogs compared to the ones carrying only the wild-type allele, both at the genome-wide and chromosome 16 level (**Figure 2**). Interestingly, North American wolves carrying the *K^B^* allele did not display a genome-wide excess of allele sharing with dogs. However, when restricting the analysis to chromosome 16, D-statistic estimates were notably elevated in North American *K^B^* carriers compared to wild-type individuals, although not reaching statistical significance (**Supplementary Figure S3**).

**Figure 2:**
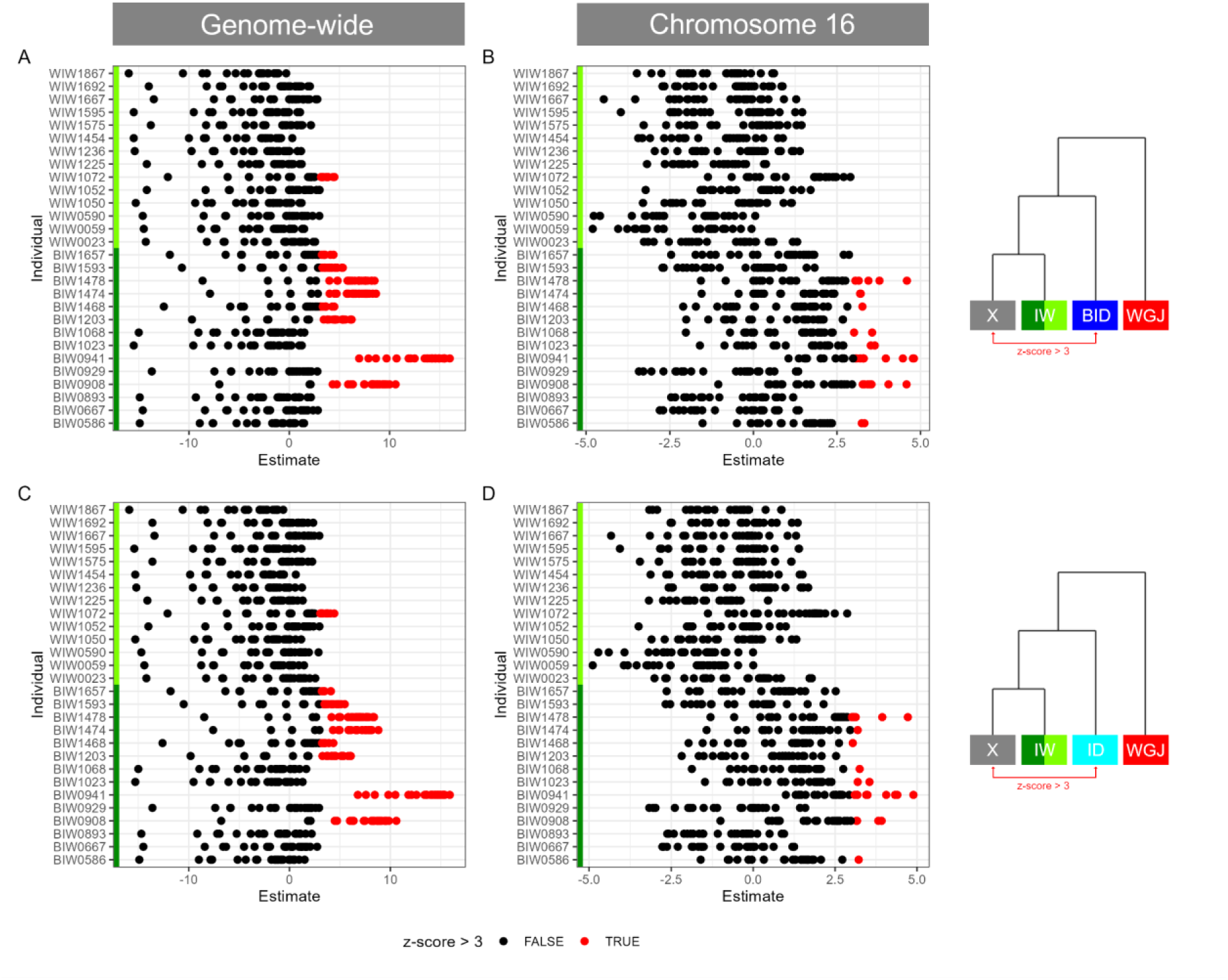
D-statistics computed (A, C) genome-wide and (B, D) on chromosome 16, testing gene flow between Italian wolves and dogs. In panel A and B, gene flow was tested between Italian wolves (BIW and WIW) and Italian dogs carrying the *K^B^* allele (BID), using golden jackals (WGJ) as the outgroup. Each Italian wolf was tested in turn (X) in pairwise comparisons with all other Italian wolves (IW) following the model: (((X, IW), BID), WGJ). In panels C and D, D-statistics were computed relative to Italian dogs carrying the wild-type *k^y^* allele (ID), using WGJ as the outgroup (((X, IW), ID), WGJ). A Z-score > 3 indicates excess allele sharing between X and dogs (BID or ID) compared to IW.

The haplotype structure around the K-locus showed two distinct haplogroups in BIW (hereafter named BIW1 and BIW2), with no variation across 3Mb on chromosomes carrying the *K^B^* allele (**Figure 3A**). Therefore, we conducted the LAI analysis separately for BIW1 and BIW2. The total length of genome-wide fragments assigned to the domestic reference was, on average, higher in BIW1 than in BIW2 (385Mb and 78.8Mb, respectively), as well as the mean number of dog segments (37 and 9, respectively).

**Figure 3:**
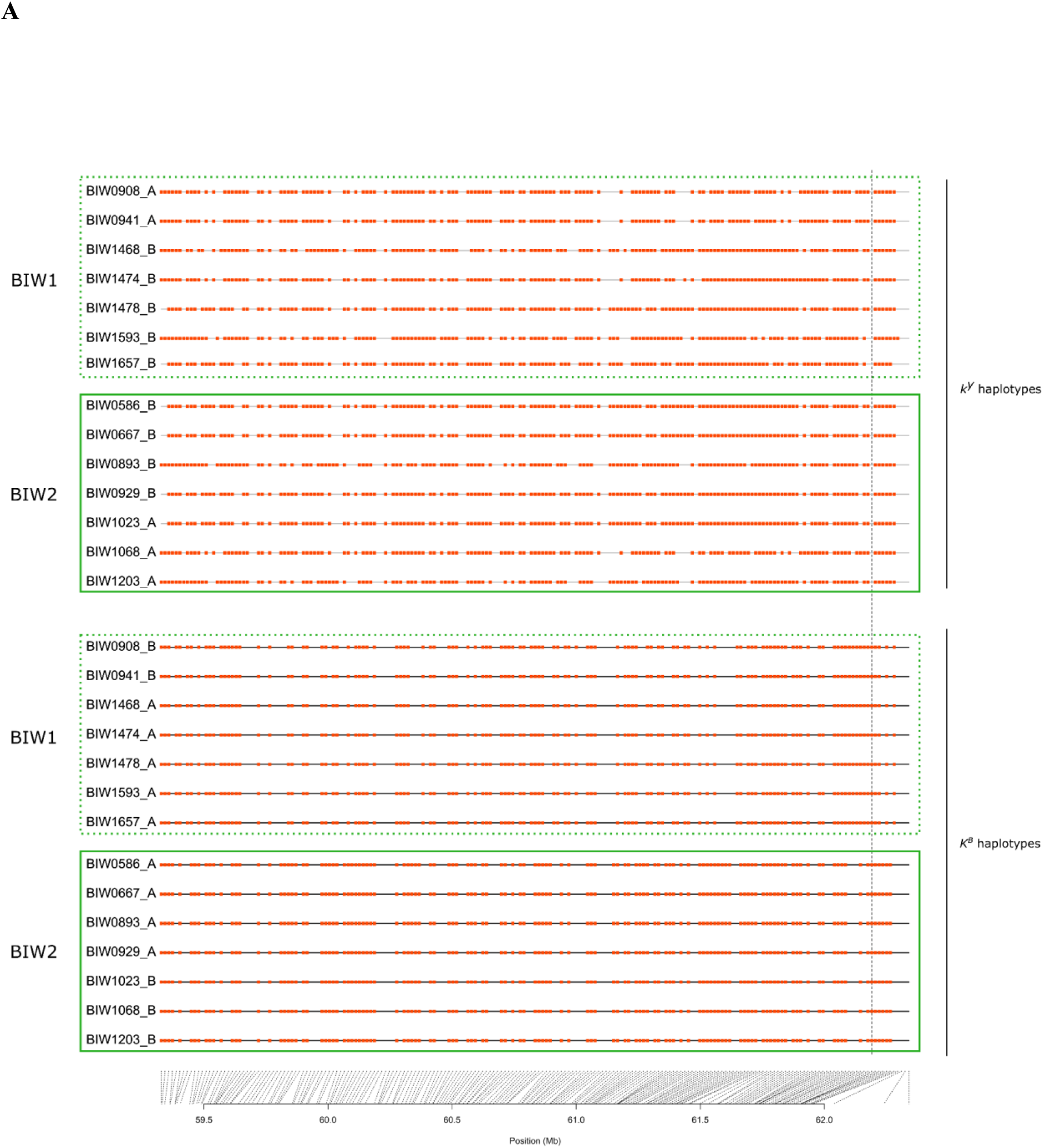
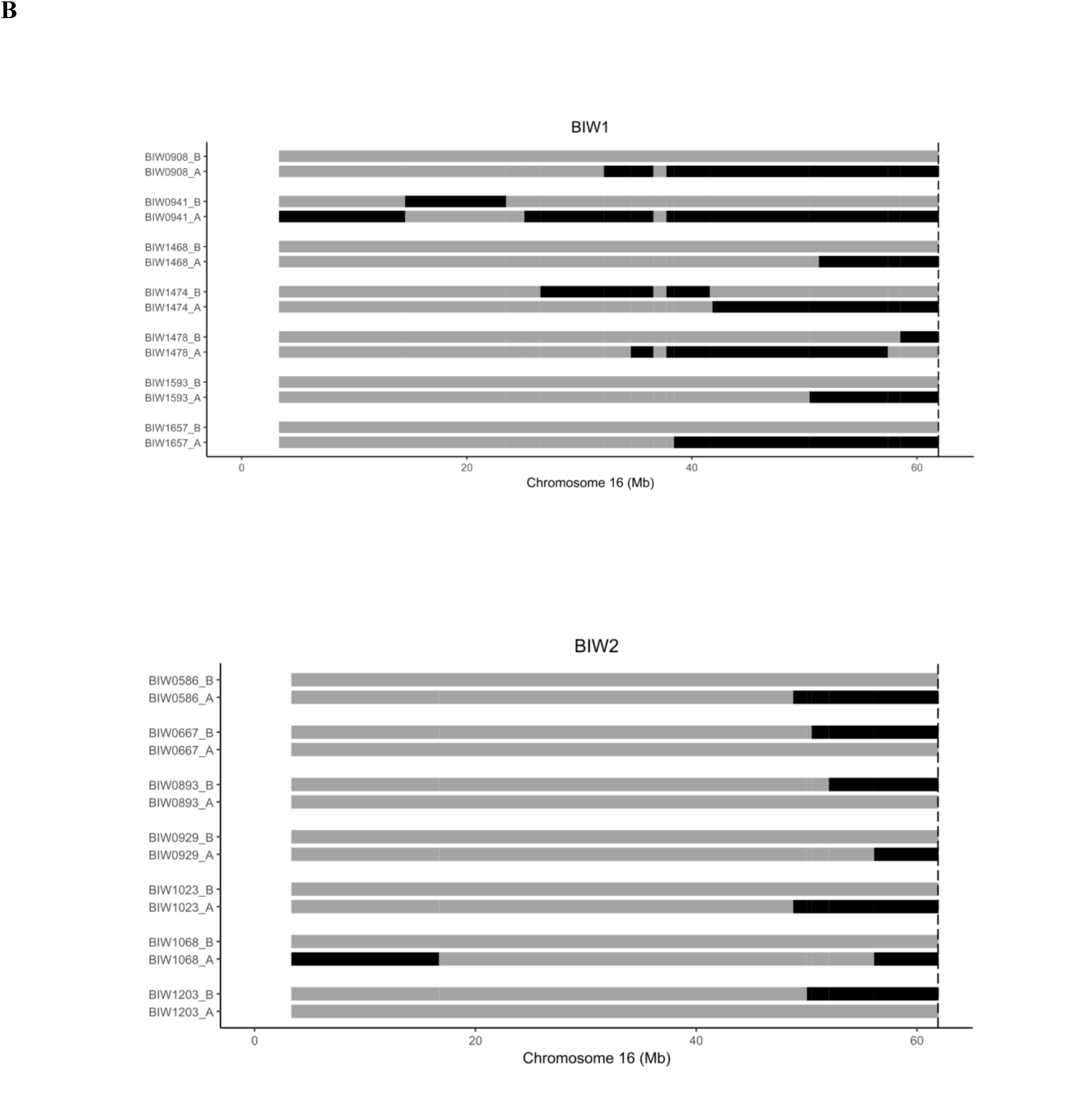
Introgression from dogs into Italian wolves. (A) Haplotype structure around the *K*^B^ allele (indicated by the vertical dashed). The chromosomes are grouped according to the allele at the K-locus: ancestral (*k*^y^) and derived (*K*^B^). The red dots represent the minor allele at each SNP. In the case of the *K^B^* allele, as its frequency was exactly 0.5, we set it as the derived allele, with *k^y^*considered ancestral. Haplotype pairs belonging to BIW1 and BIW2 haplogroups are represented within dotted and solid line rectangles, respectively. (B) Local ancestry inference along chromosome 16 for BIW1 and BIW2 haplogroups. Black = dog ancestry; grey = wolf ancestry.

Across individuals, chromosome 16 consistently carried signs of dog introgression in haplotypes from both BIW1 and BIW2 haplogroups. In fact, the LAI analysis assigned the genomic region containing the K-locus to the dog source population in each chromosome carrying the *K^B^* allele (**Figure 3B**). The average proportion of dog-derived fragments on this chromosome exceeded those observed in all the other chromosomes in all BIW individuals.

The local ancestry inference revealed that BIW1 haplotypes showed, on average, 59 ancestry switches, with genome-wide wolf ancestry proportions ranging from 80.6% to 97.2% (**Supplementary Table 2**). BIW2 exhibited fewer ancestry switches, on average 13, and consistently higher wolf ancestry proportions, ranging from 91.7% to 99.3% (**Supplementary Table 2**). For BIW1 haplotypes, the estimated time since admixture ranged from 7 to 9 generations, corresponding to about 21-27 years before individual sampling (2006-2014). This suggests that the inferred hybridization events occurred between approximately 1985 and 1993 (**Supplementary Table 2**). For BIW2 haplotypes, the estimated time since admixture ranged from 5 to 12 generations, corresponding to approximately 15-36 years before individual sampling (2000-2011), placing hybridization events between 1975 and 1992 (**Supplementary Table 2**).

### Patterns of diversity and selection around the K-locus

Allele-specific EHH computed around the *K^B^* allele, separately for the two BIW clusters, showed a slower decay on chromosomes carrying this allele, especially in BIW2 (**Figure 4**). These patterns may be suggestive of recent positive selection acting on the *K^B^* allele, although it could also reflect the recent introgression history of the melanistic deletion.

**Figure 4:**
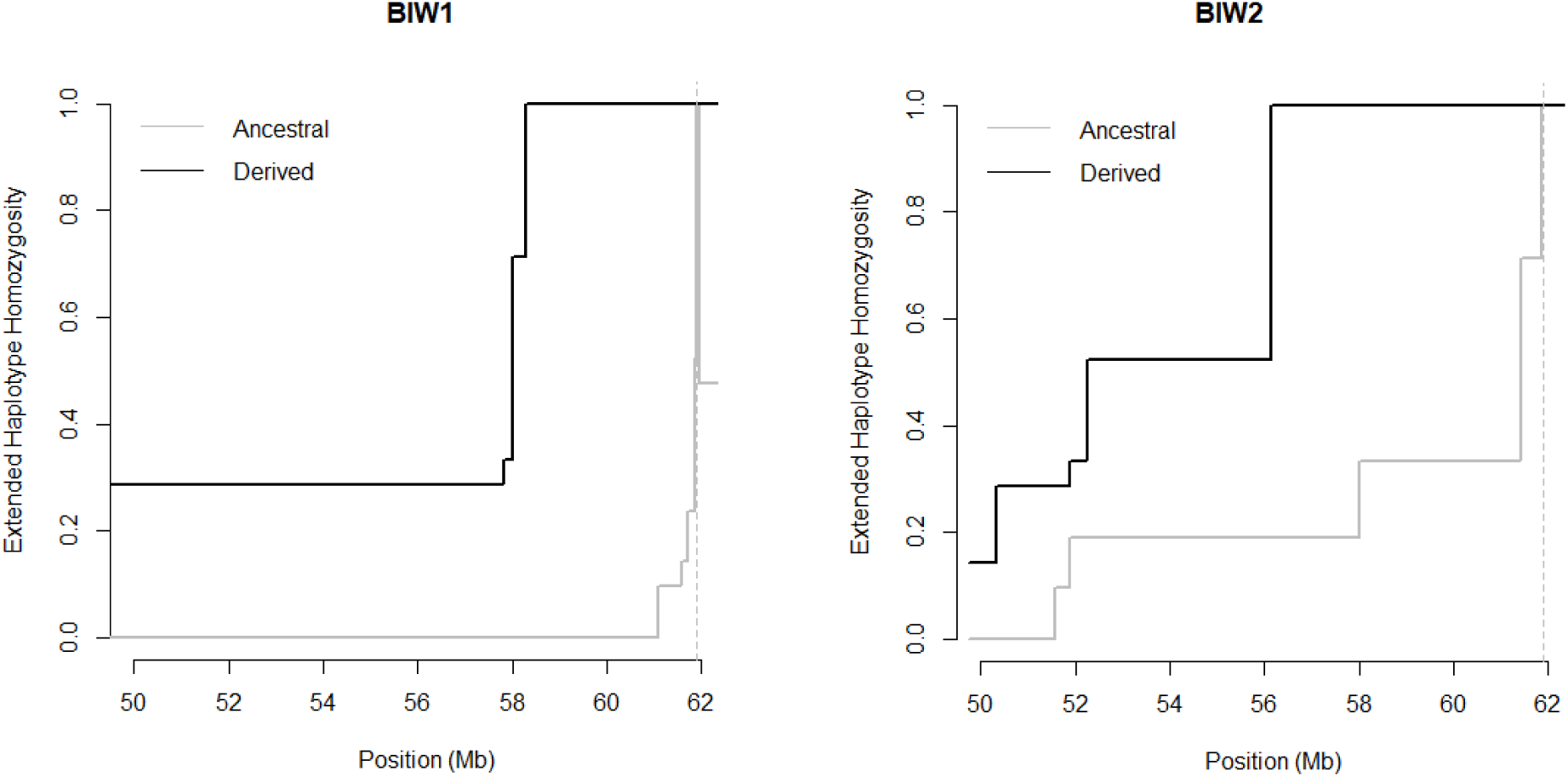
Extended haplotype homozygosity around the K-locus in BIW1 and BIW2 haplogroups. The derived allele is the *K^B^* allele, while the ancestral one is the *k^y^* allele.

Tajima’s D values, computed in non-overlapping 200-kbp windows, in BIW1, BIW2, and WIW, were slightly positive at the K-locus genomic position only in BIW2, which can be suggestive of balancing selection or population demographic contraction (**Supplementary Figure S4**).

## Discussion

In this study we analysed, through multiple ancestry reconstruction approaches, genome-wide SNP profiles of Italian wolves sampled in the northern and central Apennines, where the occurrence of black individuals has been documented since the 1970s (Boitani 1983; Apollonio et al. 2004; Anderson et al. 2009; Caniglia et al. 2013). Melanism in wolves has often been considered an indicator of hybridization with domestic dogs (Anderson et al. 2009). However, our genome-wide analyses contradict this assumption and confirm that this phenotypic trait alone cannot be considered a sufficient proxy for hybrid identification, as previously highlighted by other genetic (Caniglia et al. 2013; Roda and Philibert 2025) and genomic (Galaverni et al. 2017; Battilani et al. 2025) studies. Our multivariate principal component and maximum-likelihood assignment analyses showed that most black wolves clustered within the Italian wolf genome-wide variability, with no clear signatures of recent admixture with dogs. All 14 black wolves we analysed carried a single copy of the *K^B^* allele – known to cause the melanistic phenotypes in dogs and wolves (Anderson et al. 2009; Caniglia et al. 2013) – but only six were clearly identified as recent wolf–dog hybrids. These individuals fall within the first four backcross generations, based on their genome-wide ancestry patterns (Caniglia et al. 2020).

Allele-frequency-based tests revealed signals of domestic gene flow in a subset of BIW individuals, both genome-wide and on chromosome 16, whereas other BIW individuals exhibited non significant Z-scores comparable to those observed in wild-type wolves. This indicates that introgressed variants can persist in wolf genomes even after dilution of dog ancestry through extensive backcrossing, concordant with other genomic studies conducted on the Iberian wolf (Lobo et al. 2025; Sarabia et al. 2025), European wildcat (Mattucci et al. 2019) and European wild boar (Dzialuk et al. 2018; Fabbri et al. 2023). Interestingly, our allele-frequency-based analyses showed similar patterns of gene flow when performed using as reference dog groups with or without the *K^B^* allele. This is consistent with theoretical expectations (Martin et al. 2015), as these methods can capture genome-wide and chromosome-level ancestry patterns rather than variation at a single locus.

Phased haplotypes used for LAI analysis, led to the identification of two distinct haplogroups around the K-locus, as early hypothesized by Galaverni et al. (2017). Our LAI analyses confirmed the domestic origin of both haplogroups, which substantially differed in the total length, mean number of dog fragments, and geographic distribution. This pattern is consistent with, at least, two independent admixture events, which potentially occurred at different times and in different geographical areas. This scenario differs from that described for North American black wolves, which apparently originated from a single ancient introgression (Schweizer et al. 2018). The haplogroup with higher dog content (BIW1) was present in five recent hybrids and two non-admixed individuals, according to ADMIXTURE results. These individuals were mostly sampled in the central-western Apennines (southern Tuscany and northern Lazio regions), with a single female (BIW1468) sampled on the eastern coast of central Italy (Marche region), representing a possible disperser – or the progeny of previous dispersers – that maintained this melanistic legacy (**Figure 5**). Conversely, the haplogroup with lower dog content (BIW2) included only one recent hybrid male (BIW1203) and six non-admixed wolves all sampled in the northern-eastern Apennines (Emilia Romagna and Liguria regions; **Figure 5**).

**Figure 5:**
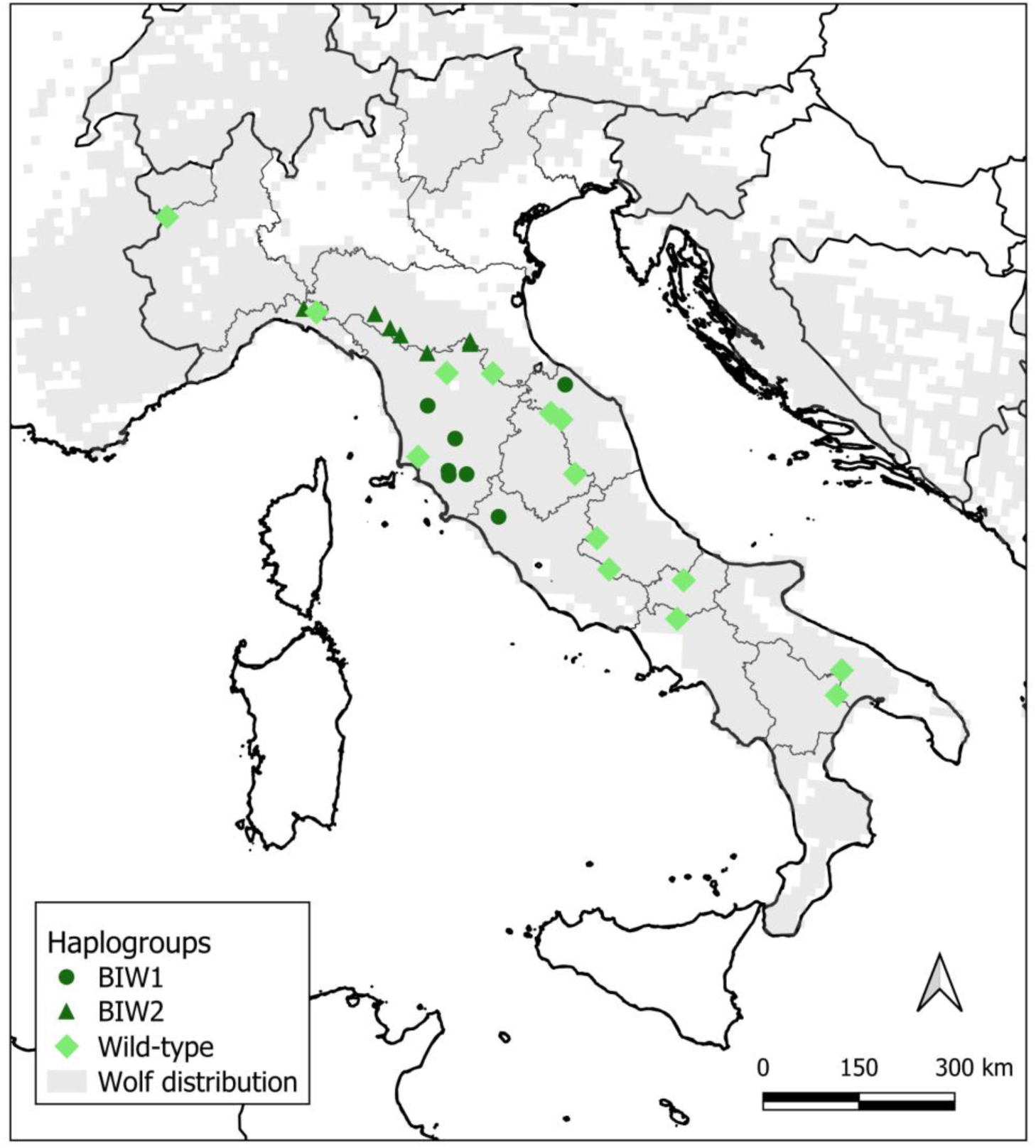
Sampling locations of the Italian wolves analyzed in this study. Wolves with the *K^B^* allele (BIW1 and BIW2, both n=7) are shown as dark green symbols, while individuals homozygous for the wild-type *k^y^* allele (WIW, n=14) are represented by light green diamonds. The wolf distribution was taken from Kaczensky and colleagues (2024).

To estimate the time since admixture in the two haplogroups, we applied a dating approach based on introgressed haplotype length (Pool et al. 2009; Maples et al. 2013), widely used for deeply introgressed populations (i.e., humans (Norris et al. 2020), canids (Wang et al. 2020), and bovids (Yan et al. 2024)), and the number of recombination switches due to backcrossing (Johnson et al. 2011). Such methods might be less performing when applied to SNPchip data than to whole-genome sequences, and the resulting outcomes should be taken with caution for possible timing underestimates due to a limited detection of shorter fragments. Nevertheless, they allowed us to infer that admixture events associated with the BIW2 haplotypes are likely older, potentially dating back to the 1970s, during the last anthropogenic population bottleneck of the Italian wolf population (Lucchini et al. 2004). In contrast, the admixture events associated with the BIW1 haplotypes might be more recent, potentially dating to the 1980s, during the population re-colonization phase along the Apennines and towards coastal areas (Galaverni et al. 2017). This is consistent with recent evidence showing that wolf–dog hybridisation tends to be more prevalent during phases of population re-expansion (Lobo et al. 2023).

Given the high dispersal capacity of wolves (Valière et al. 2003; Ciucci et al. 2009; Andersen et al. 2015), the spatial distribution of these haplogroups might reflect localized introgression events rather than a long-term population substructure (Fabbri et al. 2007; Galaverni et al. 2017).

Our allele-specific extended haplotype homozygosity (EHH) analysis revealed that the genomic region around the *K^B^* deletion showed signals compatible with recent events of introgression but also with positive selection in both the haplogroups we detected, consistent with previous findings in North American black wolves (Anderson et al. 2009; Schweizer et al. 2018). However, in North American wolves, where the *K^B^* introgression occurred approximately 7,000 years ago, recombination and purifying selection may have purged most dog-derived genomic regions, possibly leaving only few hitchhiked traces of dog ancestry surrounding the adaptive *K^B^* allele. This would explain the low genome-wide domestic ancestry observed in this population (Schweizer et al. 2018). Additionally, our allele frequency spectrum analysis based on Tajima’s D also revealed patterns compatible with balancing selection in the genomic region surrounding the *K^B^* allele in BIW2 individuals. Although positive Tajima’s D values can also arise from demographic or genetic processes, such as bottlenecks (Gattepaille et al. 2013) and gene flow (Ojeda et al. 2008), these latter cannot fully explain the results for BIW2 wolves. Indeed, both BIW1 and BIW2 individuals shared the same traces of the bottleneck that affected the entire Italian wolf population (Lucchini et al. 2004) and showed admixture patterns with dogs. Therefore, the patterns observed in BIW2 individuals are consistent with balancing selection acting on the *K^B^* haplotype. The contrasting patterns between BIW1 and BIW2 individuals might be explained by their different admixture times, which likely determined the emergence of selection signatures only in the haplogroup carrying older introgressed tracts. However, such findings are preliminary and should be taken with caution because, in contrast to other selection studies using targeted capture or whole-genome sequences (Schweizer et al. 2018; Liu et al. 2025; Yudin et al. 217; Rocha et al. 2023), the low SNP density in our dataset precluded additional testing.

Nevertheless, despite these discrepancies, both North American and Italian black wolves may exhibit possible selective advantages associated with the 3-bp deletion, representing likely cases of adaptive introgression. Consistent with evidence from melanistic North American wolves (Cubaynes et al. 2022), which showed enhanced immune performance during outbreaks of canine distemper virus (CDV), the spread of this variant in Italian wolves may be also driven by pathogen-mediated selection. Indeed, during the population re-expansion in the 2000’s, encounters with other sympatric mammals susceptible to the disease may have facilitated pathogen circulation. Ricci et al. (2021) reported that the red fox (*Vulpes vulpes*) and the domestic dog are among the main vectors for the spread of the CDV disease in central Italy. Bianco et al. (2020) also documented several outbreaks in red foxes, stone martens (*Martes foina*), and badgers (*Meles meles*) in northern Italy, suggesting that increasing carnivore density in recovering ecosystems could raise disease incidence even among wolves. Necropsy analyses of more than one hundred Italian wolf carcasses showed that, while sarcoptic mange was rather common in wild-type individuals, any of the black wolf examined exhibited apparent signs of this disease (C. Musto, personal communication). These findings suggest a possible immune resistance of *K^B^* allele carriers to CDV, which is consistent with other studies showing that additional genetic variation in loci involved in immune response are particularly prone to introgression (Fijarczyk et al. 2018; Talarico et al. 2021; Vespasiani et al. 2022). Adaptive introgression of domestic immune-related variants can, therefore, represent an important evolutionary mechanism for the long-term survival of wildlife populations. Clear examples include the beneficial introgression of domestic goat MHC alleles (Grossen et al. 2014) and immune system loci (Münger et al. 2024) in the Alpine ibex, domestic cat disease resistance genes in the European wildcat (Howard-McCombe et al 2023) and the heterozygote advantage at MHC loci described in Italian wolves (Galaverni et al. 2015).

In this context, the persistence and spread of the *K^B^*allele in wolves may reflect a complex evolutionary mechanism resulting from a trade-off between survival rates and reproductive success, potentially maintained through heterozygote advantage of the *K^B^* carriers over both homozygote genotypes (Coulson et al. 2011). Interestingly, although *K^B^/K^B^* homozygotes are present in dogs, they were almost never detected in the Italian wolves over the last 25 years (Randi et al. 2014; Caniglia et al. 2020; Gervasi et al. 2024), except for two single cases reported in peninsular Italy (Lorenzini et al. 2025). Our findings suggest that, similarly to North American wolves, *K^B^/K^B^* homozygotes are extremely rare in Italy. This might be due to a disassortative mating mechanism among melanistic individuals, as observed in North American wolves (Hedrick et al. 2016) and to a reduced fitness of *K^B^/K^B^* individuals (Coulson et al. 2011). Indeed, studies in North American wolves suggest that homozygous melanistic individuals have lower survival during the first months of life, possibly reflecting pleiotropic effects affecting immune and physiological functions (Schweizer et al. 2018; Cubaynes et al. 2022).

## Conclusions

All Italian black wolves we genomically examined carried the *K^B^*allele within dog introgressed haplotypes, similarly to North American black wolves (Anderson et al. 2009). However, the *K^B^* allele may not be the single genomic basis for black coat colour in Italian wolves, as at least two apparently black individuals without the melanistic deletion have been already reported in the Italian population (Caniglia et al. 2013). Future studies based on whole-genome data are therefore required to identify additional melanistic genetic variants associated with black coat colour in wolves.

On the other hand, it would be also important to investigate phenotypic variation within a larger sample of wolves carrying the *K^B^* allele, which might be influenced by the pleiotropic effect of other genes, including, especially, those responsible for coat colour patterns as it happens in dogs (Kaelin and Barsh 2012).

Similarly, further whole-genome studies are needed to better understand the fate of the *K^B^*allele and other dog variants which, like the *K^B^* allele, might be responsible for adaptive introgression and other microevolutionary changes in the Italian wolf population.

From a conservation perspective, recent recommendations emphasize prioritizing the identification and management of recent wolf-dog hybrids and early-generation backcrosses, which retain the highest proportion of dog ancestry (Caniglia et al. 2020; Stronen et al. 2025). Our results demonstrate that relying on a single phenotypic trait such as black coat colour is misleading, as most black wolves molecularly analysed showed limited or negligible dog ancestry. Therefore, while long-term non-invasive genetic sampling and camera-trap surveys will be crucial to track the dynamics of introgressed alleles and to understand how ecological and anthropogenic factors could shape their persistence in wild populations, hybrid detection and management decisions should primarily rely on genomic evidence.

## Data availability

The raw data used in this study will be made available after acceptance for publication on the digital repository Zenodo.

## Acknowledgements

We warmly thank all the people who contributed to the collection of the biological samples which were genotyped for this study and the personnel of Conservation Genetics Unit at ISPRA for processing the samples and leading the study.

## Ethical statements

No ethics permit was required for this study, and no animal research ethics committee prospectively was needed to approve this research or grant a formal waiver of ethics approval since the collection of wolf and jackal samples involved dead animals. Dog blood samples were collected by veterinarians during health examinations with a not-written (verbal) consent of their owners (students/National Park volunteers/or specialized technician personnel), since they were interested in wolf conservation studies and monitoring projects. Moreover, there is not a relevant local law/legislation that exempts our study from this requirement. Additionally, no anesthesia, euthanasia, or any kind of animal sacrifice was applied for this study and all blood samples were obtained aiming at minimizing the animal suffering.

