## Supplementary figures and images for "Wolves in black: multiple introgressions and natural selection may explain melanism in Italian wolves"

### Supplementary file 2

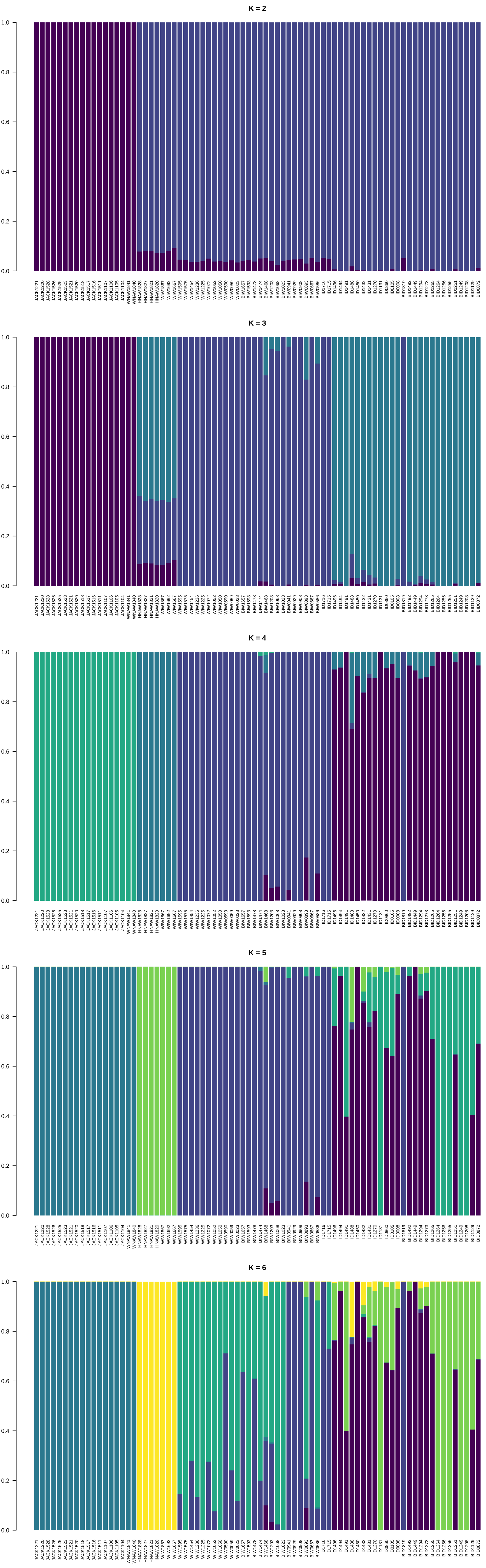
