## Supplementary Material for "Wolves in black: multiple introgressions and natural selection may explain melanism in Italian wolves"

**Supplementary table 1:** Admixture proportions of genome-wide ancestry assignment for the number of ancestral components ranging from K = 2 to K = 6. Tables provided as separate file named “Supplementary file 1.xlsx”.

**Supplementary table 2:** Summary of admixture characteristics for individuals assigned to the BIW1 and BIW2 groups. The table reports genome-wide admixture proportions attributed to wolf and dog ancestries, the number of ancestry switches between wolf and dog tracts inferred from local ancestry analysis, the estimated admixture time (in generations before sampling), the year of sampling, and the corresponding admixture time expressed in years before present.

| **BIW1** | | | | | | |
| --- | --- | --- | --- | --- | --- | --- |
| **Sample ID** | **Wolf ancestry proportion** | **Dog ancestry proportion** | **Number of switches** | **Admixture time (in generations)** | **Sampling year** | **Admixture date** |
| BIW1657 | 0.955 | 0.045 | 29 | 6.86 | 2014 | 1993 |
| BIW1593 | 0.909 | 0.091 | 71 | 8.76 | 2013 | 1987 |
| BIW1478 | 0.929 | 0.071 | 58 | 9.00 | 2012 | 1985 |
| BIW1474 | 0.924 | 0.076 | 46 | 6.68 | 2013 | 1993 |
| BIW1468 | 0.972 | 0.028 | 24 | 8.86 | 2013 | 1986 |
| BIW0941 | 0.806 | 0.194 | 112 | 7.30 | 2007 | 1985 |
| BIW0908 | 0.886 | 0.114 | 71 | 7.13 | 2006 | 1985 |
| **BIW2** | | | | | | |
| **Sample ID** | **Wolf ancestry proportion** | **Dog ancestry proportion** | **Number of switches** | **Admixture time (in generations)** | **Sampling year** | **Admixture date** |
| BIW1203 | 0.917 | 0.083 | 48 | 6.39 | 2011 | 1992 |
| BIW1068 | 0.988 | 0.012 | 13 | 11.6 | 2010 | 1975 |
| BIW1023 | 0.991 | 0.009 | 6 | 6.86 | 2008 | 1987 |
| BIW0929 | 0.991 | 0.009 | 9 | 9.84 | 2007 | 1977 |
| BIW0893 | 0.992 | 0.008 | 4 | 5.17 | 2005 | 1989 |
| BIW0667 | 0.992 | 0.008 | 5 | 6.06 | 2002 | 1984 |
| BIW0586 | 0.993 | 0.007 | 6 | 8.27 | 2000 | 1975 |

**Figure S1**: Admixture plots for K = 2 to K = 6 ancestral components. Figure provided as a separate file named “Supplementary file 2.pdf”.

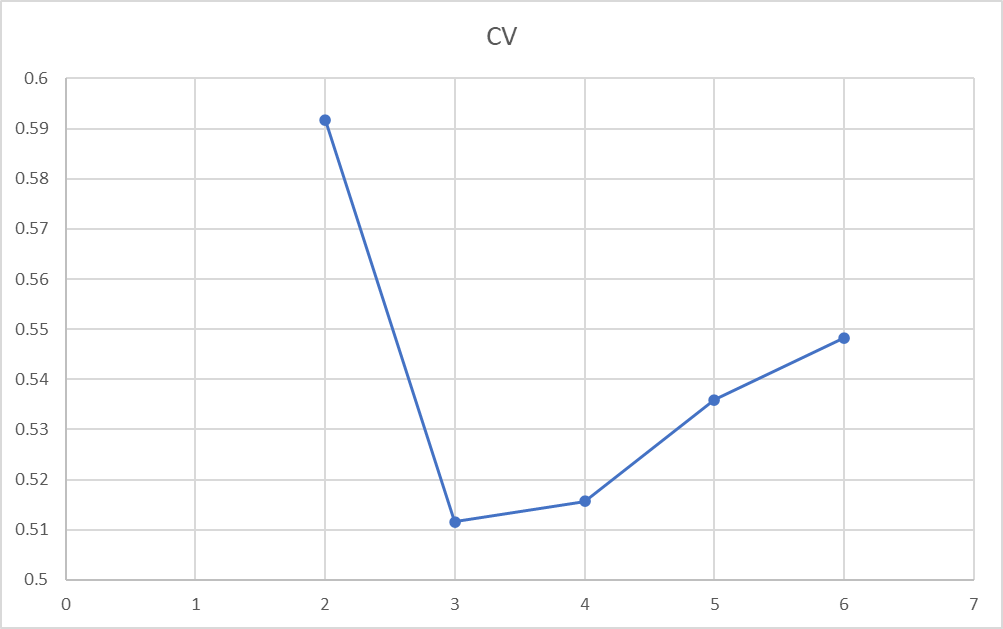

**Figure S2**: cross-validation (CV) error from ADMIXTURE for K = 2 to K = 6 applied on the genome-wide dataset.

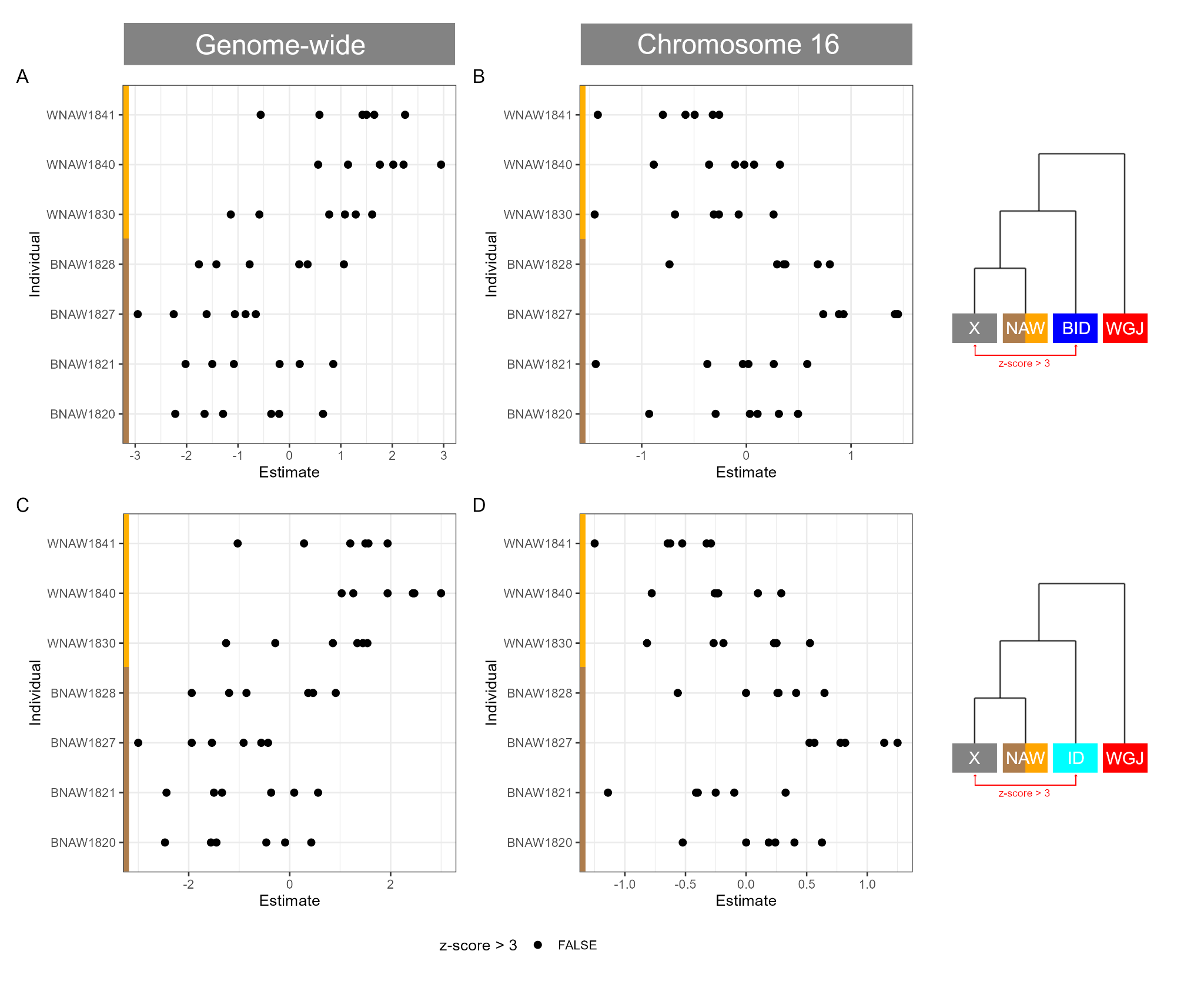

**Figure S3**: D-statistics computed (A) genome-wide and on (B) chromosome 16 testing gene flow between each North American wolf individual (X) or each other North American wolf individual (NAW), relative to Italian dogs carrying the *K^B^* deletion (BID), using golden jackals (WGJ) as the outgroup (((X, NAW), BID), WGJ). D-statistics computed (C) genome-wide and on (D) chromosome 16 gene flow between each North American wolf individual (X) or each other North American wolf individual (IW), relative to Italian dogs with the wildtype *k^y^* allele only (ID), using WGJ as the outgroup (((X, NAW), ID), WGJ). A Z-score > 3 indicates greater shared genetic drift between X and BID compared to NAW and BID (panels A and B), and X and ID compared to NAW and ID (panels C and D).

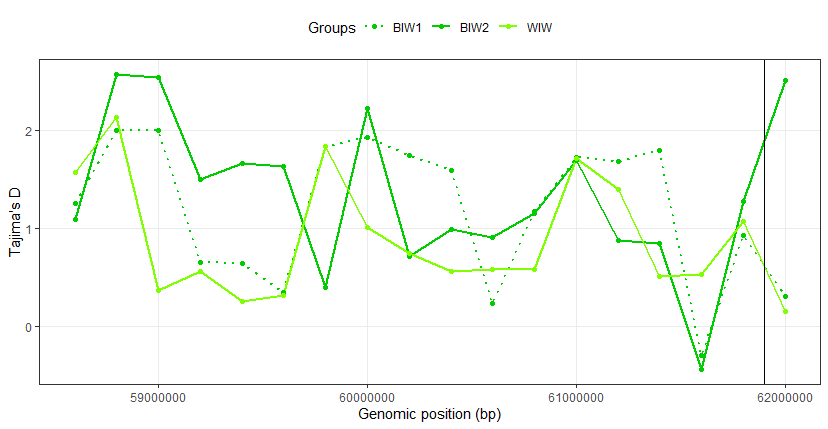

**Figure S4**: Distribution of Tajima’s D computed in non-overlapping 100 kbp windows. Only the last ~4 Mbp of chromosome 16 are shown. The position of the K-locus is indicated by the black vertical line at the end of the chromosome.
